# Generating new FANCA-deficient HNSCC cell lines by genomic editing recapitulate the cellular phenotypes of Fanconi anemia

**DOI:** 10.1101/2020.10.03.324921

**Authors:** Ricardo Errazquin, Esther Sieiro, Pilar Moreno, María José Ramirez, Corina Lorz, Jorge Peral, José Antonio Casado, Francisco J. Roman-Rodriguez, Helmut Hanenberg, Paula Río, Jordi Surralles, Carmen Segrelles, Ramon Garcia-Escudero

## Abstract

Fanconi anemia (FA) patients have an exacerbated risk of head and neck squamous cell carcinoma (HNSCC). Treatment is challenging as FA patients display enhanced toxicity to standard treatments, including radio/chemotherapy. Therefore better therapies as well as new disease models are urgently needed. We have used CRISPR/Cas9 editing tools in order to interrupt the human *FANCA* gene by the generation of insertions/deletions (indels) in exon 4 in two cancer cell lines from sporadic HNSCC having no mutation in FA-genes: CAL27 and CAL33 cells. Our approach allowed efficient editing, subsequent purification of single-cell clones, and Sanger sequencing validation at the edited locus. Clones having frameshift indels in homozygosis did not express FANCA protein and were selected for further analysis. When compared with parental CAL27 and CAL33, *FANCA*-mutant cell clones displayed a FA-phenotype as they i) are highly sensitive to DNA interstrand crosslink (ICL) agents such as mitomycin C (MMC) or cisplatin, ii) do not monoubiquitinate FANCD2 upon MMC treatment and therefore iii) do not form *FANCD2* nuclear foci, and iv) they display increased chromosome fragility and G2 arrest after diepoxybutane (DEB) treatment. These *FANCA*-mutant clones display similar growth rates as their parental cells. Interestingly, mutant cells acquire phenotypes associated with more aggressive disease, such as increased migration in wound healing assays. Therefore, CAL27 and CAL33 cells with *FANCA* mutations are phenocopies of FA-HNSCC cells.

## Introduction

Fanconi anemia (FA) is a rare, DNA repair deficiency disorder characterized by hypersensitivity to DNA interstrand crosslink (ICL) agents and chromosome instability (Bogliolo and Surralles, 2015, Niraj et al., 2019). In most cases the disease is autosomal recessive, with the exception of FANCB, which is X-linked, and mutations in the oligomerization domain of RAD51/FANCR, which are dominant negative. FA patients display varying degrees of developmental abnormalities, bone marrow failure (BMF) and increased cancer incidence. FA is due to functional inactivation of any one of 23 FA genes involved in DNA repair. The FA proteins together with FA-associated proteins interact in a pathway to repair ICL known as the FA pathway or the FA-BRCA pathway (Niraj et al., 2019). The pathway involves detection of the DNA crosslink at the stalled replication fork, unhooking of the crosslink, local generation of a double-strand break and the use of homologous recombination (HR) proteins downstream to repair the break. The management of the BMF disease has remarkably improved over the last 20 years predominantly due to improved outcome/survival of allogeneic hematologic stem cell transplantation, and more recently to hematopoietic gene therapy (Rio et al., 2019). Therefore, FA patient survival has increased from less than 20 years of age in the 1990s to more than 30 years observed today (Risitano et al., 2016, Alter et al., 2018). However, one of the most challenging health issues in older/transplanted FA patients is appearance of solid tumors such as head and neck squamous cell cancer (HNSCC) relatively early in life. Although first chemoprevention studies with quercetin or metformin (Li et al., 2012, Zhang et al., 2016) are currently conducted, treatment options are limited to curative surgery and some radiotherapy, as most FA patients display high toxicities to treatment with DNA alkylating agents and especially platinum derivatives, which form the backbone of solid cancer therapy. Compared with non-FA patients, FA-HNSCC is diagnosed at much advanced stages, and recurrence/metastatic disease is more frequent (Kutler et al., 2016). As a consequence, patient survival is very poor (25% at 5 years) (Kutler et al., 2016) and therefore new therapeutic options are needed.

Discovery and preclinical testing of new cancer treatments for FA patients is also dependent on the availability of clinical specimens for basic research purposes, which are rather scarce in the case of rare diseases such as FA. As only a handful of HNSCC cell lines derived from FA patients exists worldwide, and the characteristics of the cell lines including the genetic information and the sensitivity profiles to anticancer compounds are largely unknown (Montanuy et al., 2020, van Harten et al., 2019, van Zeeburg et al., 2005), we propose in this report to use genetic engineering on well-established HNSCC cell lines in order to generate new FA-deficient cell line models for studying FA-HNSCC. The main advantages of using such engineered FA cell lines are that their molecular aberrations (Ghandi et al., 2019) (Martin et al., 2014) and sensitivity profiles to many compounds have already been determined (Barretina et al., 2012) and in vivo xenograft models established (Palacios-Garcia et al., 2019). To this end, biallelic inactivation of *FANCA*, the gene most frequently mutated in FA patients (60%), was achieved by CRISPR/Cas9 editing of CAL27 and CAL33 non-FA two well-known oral cancer cell lines. The results presented here showed that edited clones display all FA-associated phenotypes, validating them as reliable FA-HNSCC models.

## Results

### Biallelic mutation in the *FANCA* gene by CRISPR/Cas9 editing in non-FA HNSCC cell lines

Most HNSCC in FA patients are localized in the oral cavity, mainly in the tongue. Therefore, we selected oral cancer cell lines from non-FA patients having no mutations in any of the 23 FA genes. Therefore, we mined into the CCLE data in the cBioPortal (https://www.cbioportal.org) and selected CAL27 and CAL33 cells, which display mean levels of the FANCA mRNA when compared with similar cell lines, and no amplification into the *FANCA* locus (Fig S1 and Table S1). CAL27 and CAL33 cells were nucleofected with a ribonucleoprotein complex constituted by the CRISPR guide RNA (gRNA) gGM10 targeting *FANCA* gene and purified Cas9 protein (see M&M) (Fig. 1A). Nucleofection conditions were setup to favor non-homologous end joining (NHEJ) repair after DNA excision by the gRNA/Cas9 complex. Before single cells were cloned and expanded, an estimation of editing efficiency was performed using a nuclease assay and specific primers (Table S2). Sanger sequencing was performed on selected clones to assess for inactivating mutations in both alleles. A high proportion (>90%) of sequenced clones from both CAL27 and CAL33 displayed *FANCA* gene editing (Fig. 1B). Knock-out score (KO-score) was estimated for each clone upon Sanger sequencing using the ICE algorithm (Hsiau et al., 2019) (Fig. 1C). Four clones of each parental cell line having highest KO-scores displayed negligible levels of FANCA protein, validating the editing approach (Fig. 1D). Two of these clones were selected for further analyses: clones c34 and c47 for CAL27, and clones c5 and c11 for CAL33. The edited sequences at the *FANCA* locus in each of these 4 clones are showed in Figure S2 and S3.

**Figure 1.**
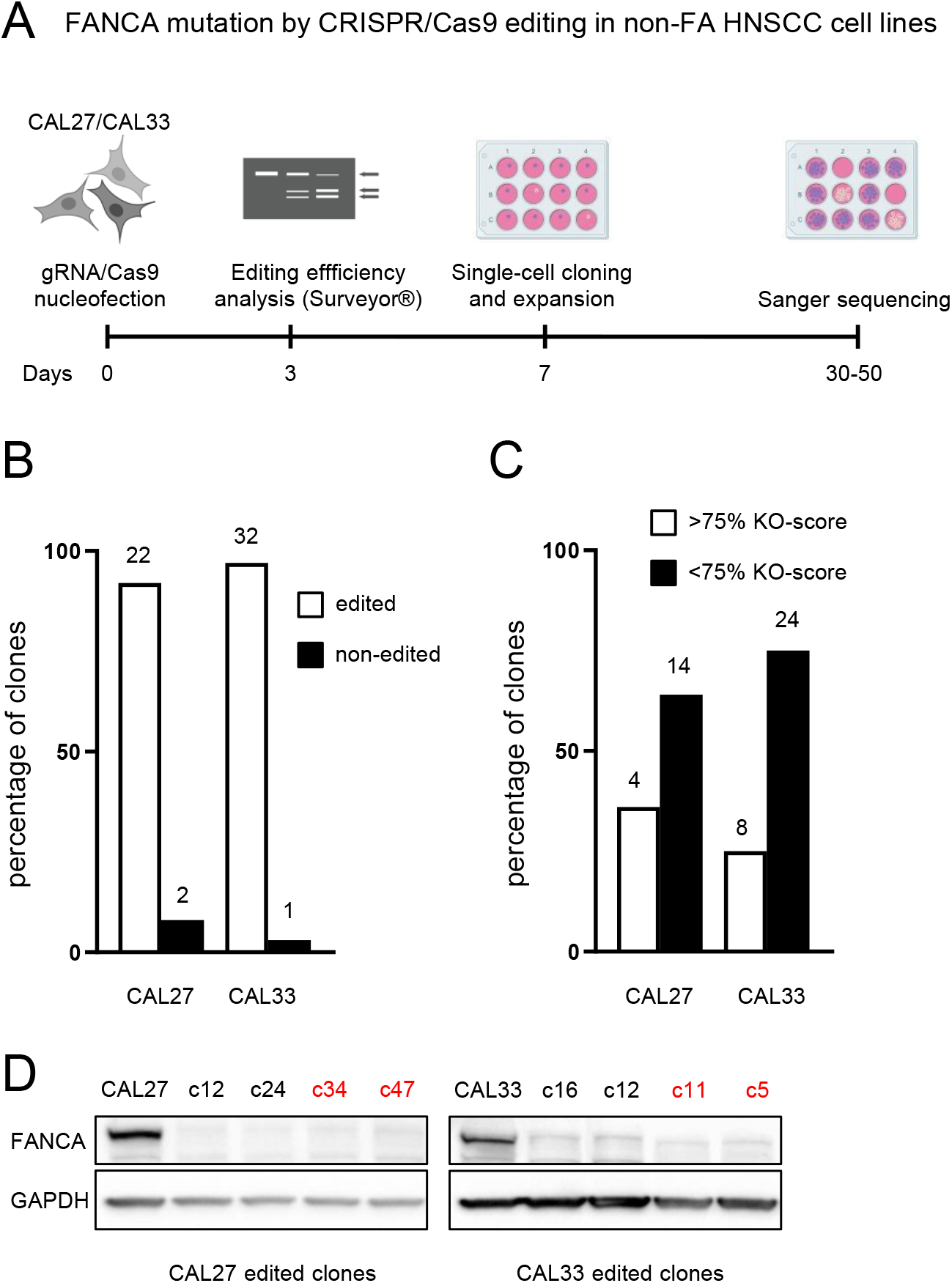
Biallelic mutation in *FANCA* gene by CRISPR/Cas9 editing in non-FA HNSCC cell lines. **A)** Schematic model of *FANCA*-mutant clones generation. CAL27 and CAL33 cells were nucleofected with a ribonucleoprotein mix of a *FANCA* gene CRISPR guide RNA (gRNA) and purified Cas9 protein. Editing efficiency was evaluated three days after by nuclease assay using the Surveyor® Mutation Detection Kit (IDT). Single cells were plated using FACS and expanded in conditioned medium. Sanger sequencing was performed on randomly selected, single-cell clones to assess for inactivating mutations in both alleles. **B)** The vast majority of sequenced clones displayed *FANCA*-gene editing, both from CAL27 and CAL33 cells. Numbers of edited or non-edited clones are shown above each bar. **C)** Knock-out score (KO-score) was estimated for each clone upon Sanger sequencing using the ICE algorithm (Hsiau et al., 2019). Between 24% and 36% of edited clones displayed high KO-scores (>75%). **D)** Clones having highest KO-scores display negligible levels of FANCA protein, validating the editing approach. Clones highlighted in red were selected for further analyses.

### *FANCA*-mutant clones are hypersensitive to ICL-agents

The hypersensitivity to the chromosome-breaking effects of ICL-inducing agents provides a reliable cellular marker for the diagnosis of FA (Giampietro et al., 1993, Auerbach, 1993). Cells from FA patients are extremely sensitive to such agents such as mitomycin C (MMC), a drug that is currently used as a diagnostic tool. Therefore, we tested the sensitivity of *FANCA*-mutant clones to MMC using cell viability assays, and the concentrations of MMC corresponding to its 50 inhibitory concentration (IC50) were calculated. IC50 values of mutant clones were 4.9 to 13.2 times lower than their corresponding, CAL27 and CAL33 parental cells (Fig 2A and Table 1). As expected, similar results were obtained with VU1365 cells, a FA patient-derived HNSCC cell line (*FANCA* mutated) previously described (van Zeeburg et al., 2005). Defective VU1365 cells were 7.4 times more sensitive to MMC than *FANCA*-complemented cells (VU1365-FANCA) obtained upon retroviral transduction of a functional *FANCA* gene. Same findings were obtained with cisplatin, another ICL-inducing agent currently used as main chemotherapy in HNSCC (Fig. 2B and Table1). Overall, sensitivity assays demonstrated that engineered *FANCA* mutated clones from non-FA HNSCC cell lines are hypersensitive to ICL-agents.

**Figure 2.**
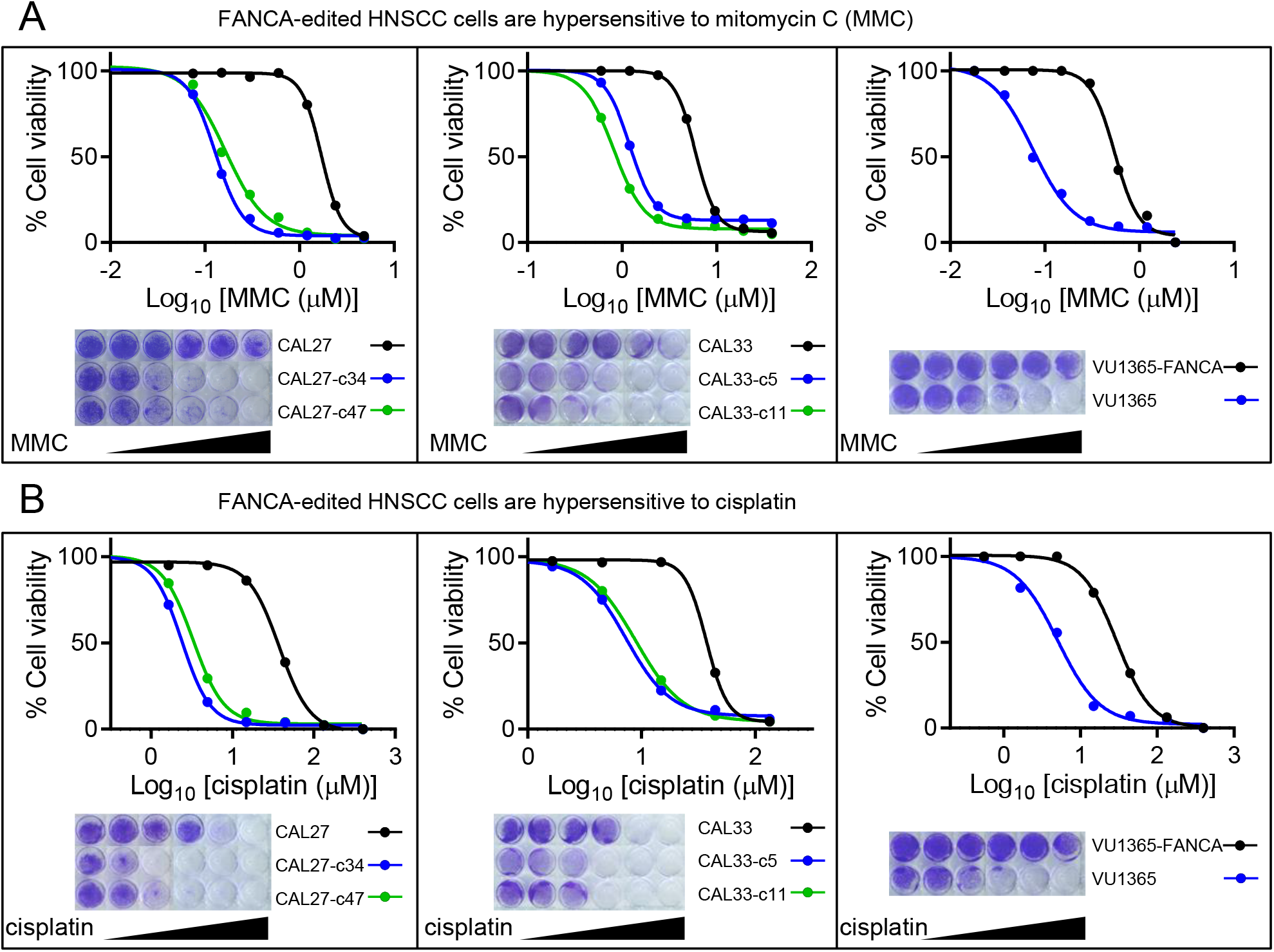
*FANCA*-edited HNSCC cells are hypersensitive to ICL agents. Cells were treated with increasing concentrations of mitomycin C (MMC) **(A)** or cisplatin **(B)** for 1 hour and grown until control cells reached the maximum confluence (5 days). Cells were stained with crystal violet, eluted with acetic acid, and color intensity was measured with absorbance at 620 nm. Data represent means ± SEMs from three different experiments for each cell line. *FANCA*-mutant CAL27 and CAL33 edited clones are more sensitive than parental cells to MMC and cisplatin insults. As expected, VU1365 cancer cells from a FA patient are also more sensitive than FANCA-complemented, VU1365-FANCA cells.

**Table 1.**
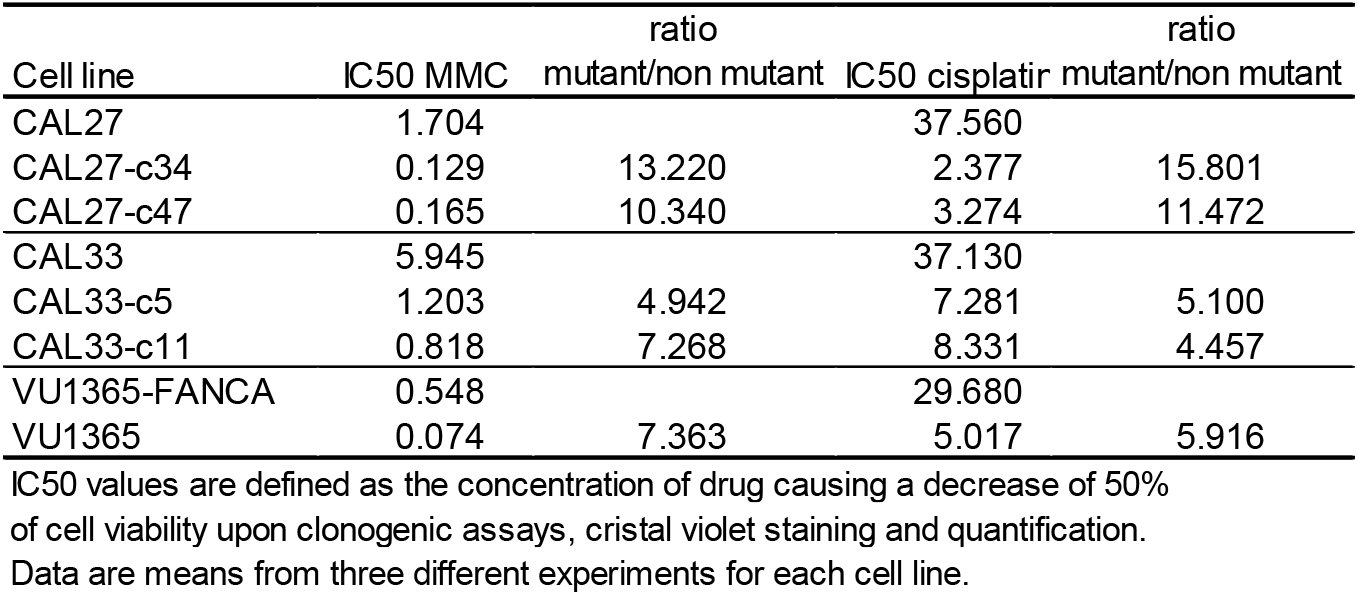
50 inhibitory concentration (IC50) values for mitomycin (MMC) and cisplatin (microM)

### *FANCA*-mutant clones are defective in FANCD2 monoubiquitination and nuclear foci formation

Early upon ICL, the FA core complex monoubiquitinates the FANCI-FANCD2 (ID2) complex which accumulates in nuclear foci (Garcia-Higuera et al., 2001). CAL27 and CAL33 cells were treated for 1 hour at IC50 concentration of MMC. As expected, a protein band of higher molecular weight over FANCD2 protein was detected, representing the monoubiquitinated form of FANCD2 (Fig 3A). As reported, a similar result was obtained with VU1365-FANCA cells (Lombardi et al., 2015). Moreover, immunofluorescence images showed that FANCD2 accumulated in nuclear foci of treated cells, corroborating that parental CAL27 and CAL33 cells have a functional FA pathway, as the VU1365-FANCA cells (Fig. 3B and C). Interestingly, CAL27-c34 and CAL33-c11 clones did not ubiquitinate FANCD2 (Fig. 3A) neither showed nuclear foci upon DNA damage (Fig. 3B and C), suggesting a defective FA pathway similar to FA-HNSCC cells VU1365 (Fig 3 C and B) (Lombardi et al., 2015) or blood cells and fibroblasts from FA patients.

**Figure 3.**
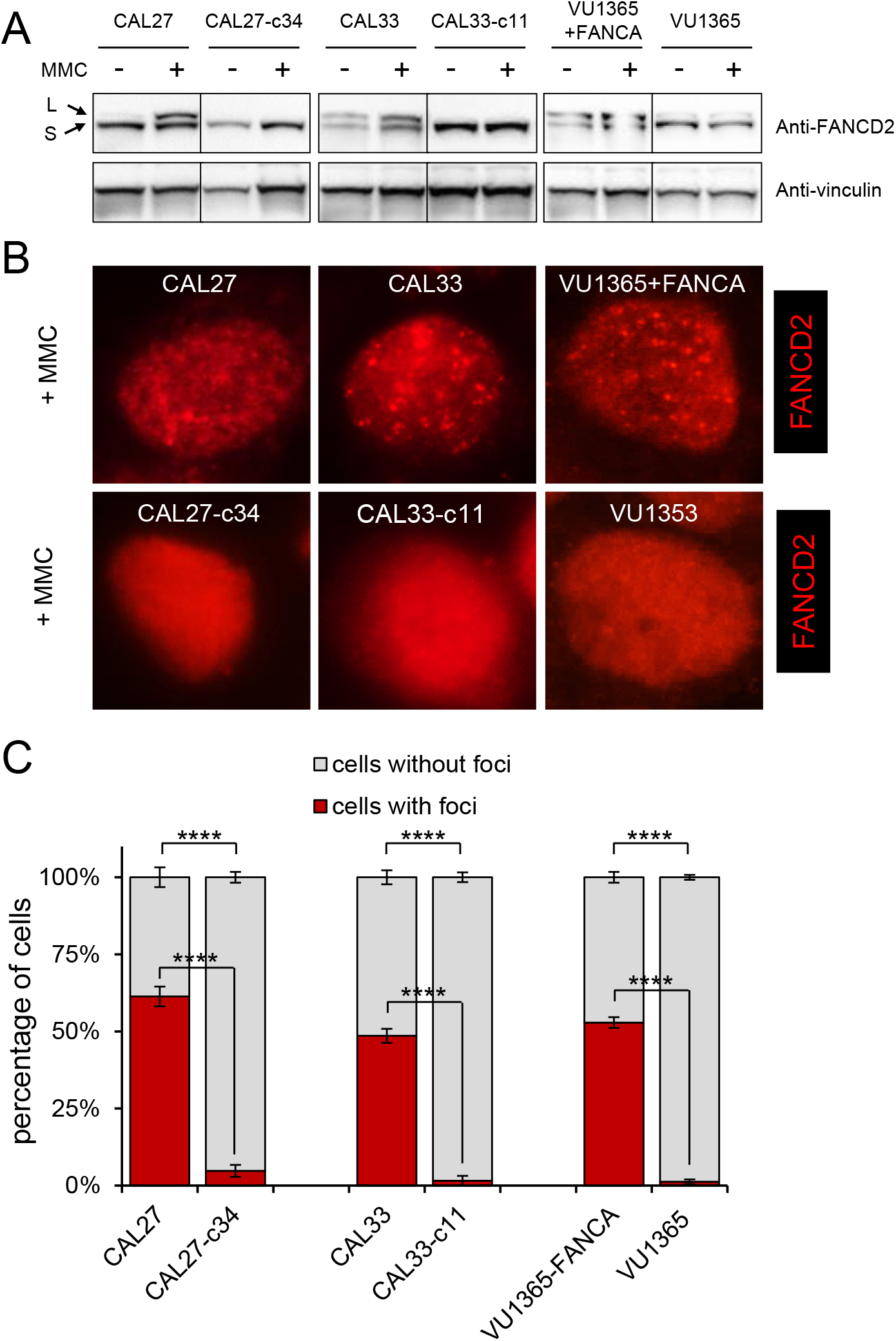
*FANCA*-edited HNSCC cells are defective in FANCD2 monoubiquitination and nuclear foci formation. CAL27, CAL33 and VU1365 parental and derivative cells were treated with IC50 concentration of MMC for 1 hour and harvested. **A)** Western blot with anti-FANCD2 antibody showing that FANCD2 monoubiquitination was not observed in *FANCA* mutant CAL27 and CAL33 clones (CAL27-c34 and CAL33-c11, respectively) suggesting a defective Fanconi pathway. As reported, parental VU1365 also lacked FANCD2 ubiquitination, which was corrected upon FANCA complementation (VU1365-FANCA). A vinculin antibody was used as housekeeping control. **B)** Immunofluorescence analysis with FANCD2 antibody. *FANCA*-mutant cells (CAL27-c34, CAL33-c11) cannot form nuclear foci after 1 hour of MMC treatment at IC50 concentration. **C)** Quantification of nuclear foci in treated and untreated cells. Data represent mean ± SEM. p-values were calculated using a two-way ANOVA test, ****: p-value ≤ 0.0001.

### Increased chromosome fragility and G2 arrest in *FANCA*-mutant clones

Micronuclei (MN) are chromosome fragments that are left behind in anaphase and appear in daughter cells as small additional nuclei. MN formation is considered a surrogate marker of chromosomal fragility. Increased MN frequency has been found in buccal mucosa cells from patients with defects in DNA-repair pathways, and has been proposed as a biomarker of cancer risk in FA patients (Ramirez et al., 2020). *FANCA*-mutant and parental CAL27 and CAL33 cells were treated with diepoxybutane (DEB) during 48 hours and the MN frequency mas measured. Results showed that mutant clones had increased MN frequency when compared with their corresponding parental cells (Fig. 4A) consistent with exacerbated chromosome fragility. Moreover, mutant cells displayed increased G2 arrest after DEB treatment (Fig. 4B), another hallmark of FA deficiency (Chandra et al., 2005). Both results support mutant clones as adequate models of FA-HNSCC disease.

**Figure 4.**
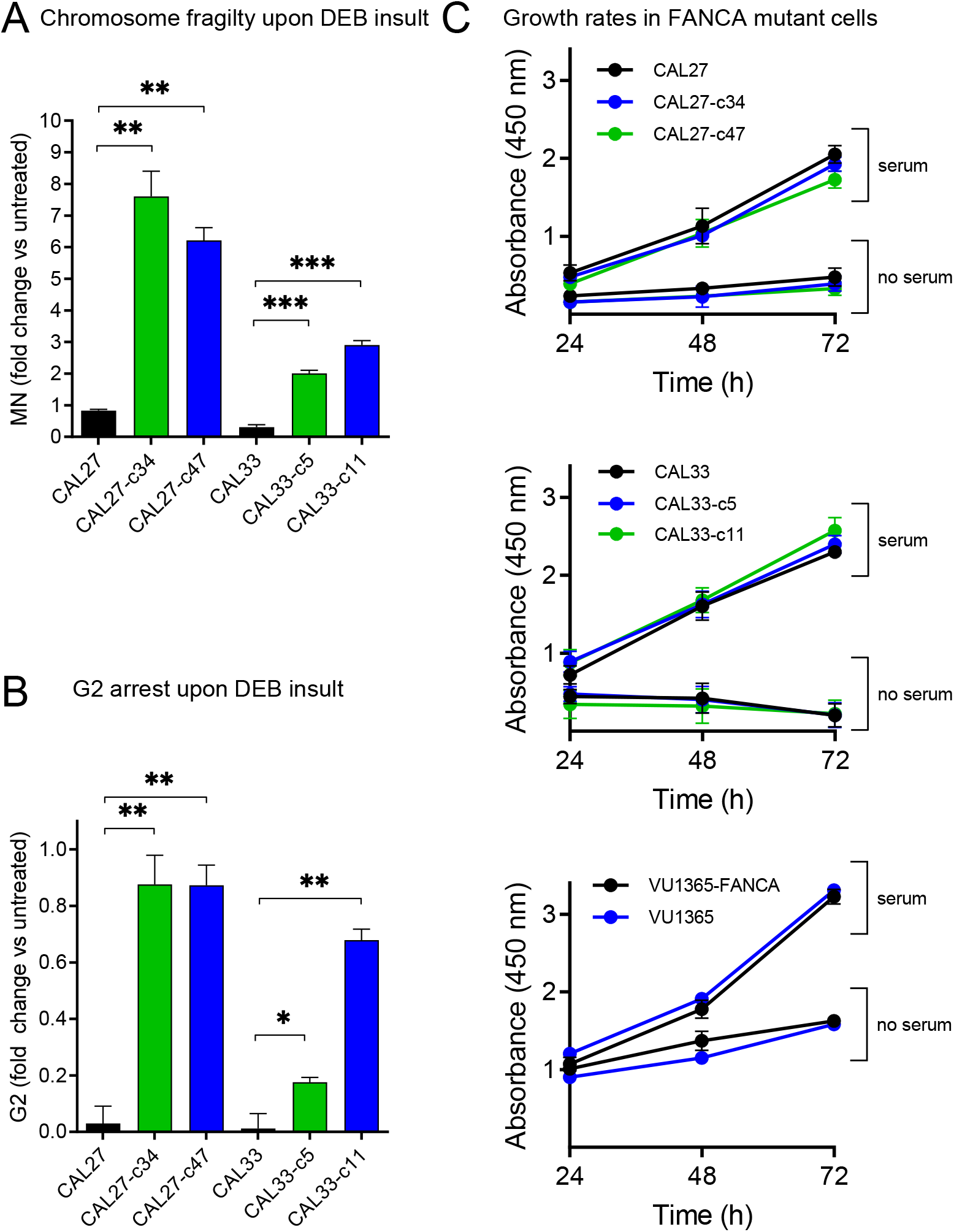
**A)** Chromosome fragility analyzed by the micronucleus (MN) test. Cells were treated with diepoxybutane (DEB), kept in culture at least one population doubling and MN were quantified with flow cytometry. FANCA-edited clones displayed increased MN formation and therefore chromosome fragility. Data are means ± SEMs from three independent experiments, each one in duplicate. **B)** Cells in G2 phase of the cell cycle were quantified upon DEB insult. *FANCA-* edited clones arrested in G2, as expected for dysfunctional FA pathway. Data are means ± SEMs from three independent experiments, each one in duplicate. **C)** HNSCC cells were seeded and maintained in culture with (serum) or without (no serum) 10% FBS for 24, 48 and 72 hours. Cell viability was measured using XTT assay and absorbance at 450 nm. *FANCA* mutation does not affect growth rate in HNSCC cells. p-values were calculated upon a T-Student test for independent samples (**A** and **B**) or upon a two-way ANOVA test (**C**). *: p-value≤ 0.05; **: p-value≤ 0.01; ***: p-value≤ 0.001; ****: p-value ≤ 0.0001.

### *FANCA*-mutation does not affect growth rates in HNSCC cells

FA genes can affect cellular growth as their function is to assure DNA integrity. Transduction with therapeutic lentiviral vectors allow proliferative advantage of corrected versus FANCA mutant, hematopoietic cells (Rio et al., 2017). Also, adult Fanconi mice (Fanca^-/-^ and Fancg^-/-^) have reduced proliferation of neural progenitor cells (Sii-Felice et al., 2008). We tested whether HNSCC cells with *FANCA*-mutation proliferate differently, both in standard (10%) or in non-serum conditions. Strikingly, results showed no differences between mutant clones and parental CAL27 and CAL33 cells, or between VU1365 and complemented VU1365-FANCA cells (Fig. 4C). Therefore, a functional *FANCA* gene does not provide proliferative advantage on HNSCC cell lines.

### *FANCA*-mutation augments cell migration in HNSCC cells

FA patients with HNSCC have poorer clinical outcomes than non-FA patients, probably due to fewer treatment options. However we cannot discard that FA deficiency might accelerate cancer progression by molecular mechanisms not fully understood. Increased *in vitro* migration/invasion was reported for sporadic HNSCC cell lines whereby FA genes were knocked-down with shRNA (Romick-Rosendale et al., 2016). We evaluated migration capacity using scratch wound healing assays in CAL27/CAL33 mutant clones and VU1365 cells. Wound closure was measured for a period of 72 hours, showing that HNSCC cells with *FANCA* mutation migrate faster and close the wound before (Fig. 5). This result suggests that HNSCC with defective FA pathway might be more aggressive.

**Figure 5.**
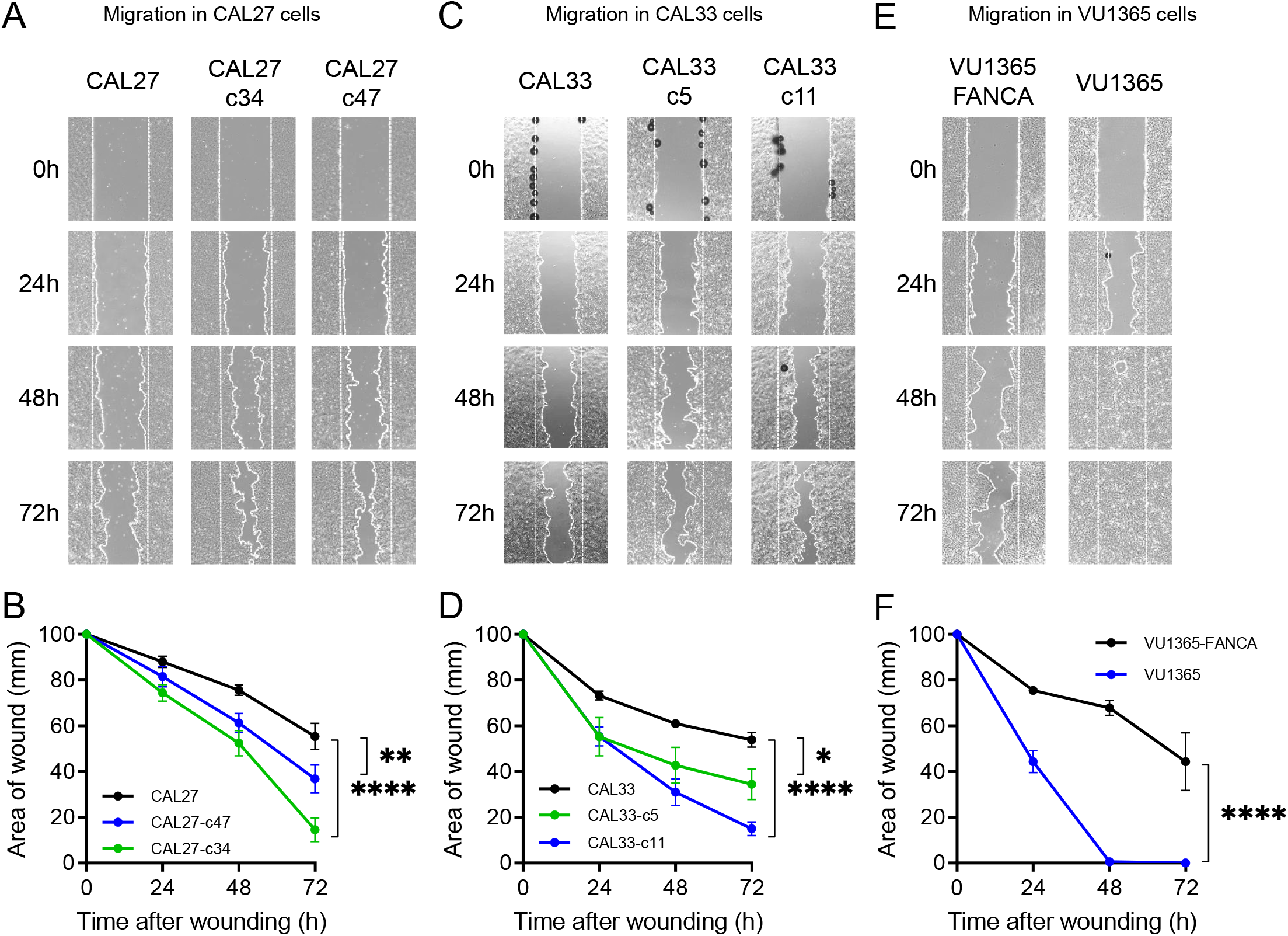
Migration analysis of *FANCA*-edited cells. **A, C** and **E**: Representative images of keratinocyte sheets that were scratch wounded to assess migration. Scale bar: 100 μm. **B**, **D** and **F**: Wound area analyzed at different times after wounding at 1% FBS. Data are mean ± SEM of one experiment with four images per time-point. p-values were calculated upon a two-way ANOVA test. *: p-value≤ 0.05; **: p-value≤ 0.01; ***: p-value≤ 0.001; ****: p-value ≤ 0.0001. CAL27 and CAL33 edited clones as well as VU1365 cells migrate faster as they closed the wounds earlier.

Overall, our CRISPR/Cas9 strategy to obtain *FANCA*-mutant clones from non-FA HNSCC cell lines gave adequate FA-HNSCC disease models as mutant clones i) are hypersensitive to ICL-agents MMC and cisplatin, ii) cannot monoubiquitinate FANCD2 and form nuclear foci upon MMC treatment, and iii) display increased chromosome fragility and G2 arrest when treated with DEB. Importantly, mutant cells migrate faster and constitute a useful tool to investigate molecular and cellular mechanisms linking the FA pathway with invasive phenotypes.

## Discussion

A few HNSCC cell lines from FA patients exist, including VU1365 which we use here. Increasing the number of cell lines, and incorporating those having classical HNSCC mutations, are necessary to find adequate new treatments for FA patients with HNSCC. Such treatments should not induce chromosome fragility to avoid toxicities that might preclude their clinical use. Therefore, we predict that our *FANCA*-mutant clones from CAL27 and CAL33 will be useful in this context.

CAL27 and CAL33 are well studied cell lines that have known aberrations/mutations in classical HNSCC genes such as *TP53* (CAL27/CAL33), *PIK3CA* (CAL33), *FAT1* (CAL27/CAL33), *CASP8* (CAL33), etc (Martin et al., 2014). Oncogenic signaling and response to cetuximab, cisplatin, and radiotherapy has been thoughtfully investigated in these cell lines (Wang et al., 2014) (Huang et al., 2016) (Eke et al., 2012). In addition, sensitivity profiles for many compounds and their association with molecular aberrations have been determined for CAL27 and CAL33 as part of the cell lines within consortiums such as CCLE (Barretina et al., 2012), Genomics of Drug Sensitivity in Cancer (Garnett et al., 2012), and Cancer Dependency Map (https://depmap.org/portal/depmap/). Therefore, links between distinct pharmacologic vulnerabilities to genomic patterns and translation into cancer patient stratification is possible for CAL27 and CAL33 (Ghandi et al., 2019). How some of these vulnerabilities change upon *FANCA* mutation should be investigated in the future.

The CRISPR/Cas9 strategy that we use results in very high editing frequencies (>90%, see Fig. 1). Possible factors include the absence of a functional p53 pathway in CAL27 and CAL33, which it has been shown to inhibit CRISPR/Cas9 editing. Also, given that mutating *FANCA* in these cells did not reduce growth rates (Fig. 4C), no negative selection of mutant versus non-mutant clones soon after nucleofection occurs. We predict that other gene editing attempts in additional FA genes should also be feasible in this setting.

CAL27 and CAL33 cell lines monoubiquitinate FANCD2 and form FANCD2 nuclear foci after ICL damage, demonstrating a functional FA pathway (Fig 3). Such activities were impeded upon biallelic inactivating *FANCA* mutations in cell clones obtained after CRISPR/Cas9 editing which also make cells hypersensitive to MMC and cisplatin (Fig. 2). Furthermore, *FANCA*-mutated clones from CAL27 and CAL33 treated with DEB displayed chromosome fragility and G2 arrest (Fig. 3A and B). All these findings clearly demonstrated that CAL27-c34 and -c47 clones, and CAL33-c5 and -c11 clones have defective FA pathway, and therefore, they constitute new FA-HNSCC cellular models.

A functional *FANCA* gene does not provide proliferative advantage to CAL27, CAL33 or VU1365-FANCA cells (Fig 3C) in cell culture. A similar result was reported for VU1365 parental and FANCA complemented cells grown in three-dimensional cultures (Lombardi et al., 2015). These results are in contrast with the reduced viability of FA-deficient hematopoietic cells. Although a synchrony between FA gene expression and the cell cycle in HNSCC cell lines has been shown (Hoskins et al., 2008), further analysis might help to understand whether the FA pathway can have role in the proliferative potential in HNSCC.

Response to current therapies of FA patients with HNSCC is very poor, partially due to toxicities to chemo/radio therapies. Moreover, most patients are diagnosed in advanced stages and recurrent/metastatic disease is frequent. However, we cannot discard an involvement of the FA pathway in tumor progression. We have found that *FANCA* mutant HNSCC cells, either from FA or non-FA patients, display increased *in vitro* migration speed when compared with FA pathway proficient counterparts (Fig. 5). This result is in line with reports indicating that HNSCC with DNA repair defects display features of invasive disease (Bakhoum et al., 2018, Bhide et al., 2016, Essers et al., 2019). More specifically, increased *in vitro* invasiveness of sporadic HNSCC cell lines where FA genes have been knocked-down with shRNA was also reported (Romick-Rosendale et al., 2016). We propose that *FANCA*-mutant clones from CAL27 and CAL33 are good tools to deepen into molecular mechanisms of tumor progression mediated by FA genes.

HNSCC is a life-threating health complication in FA patients for whom standard therapies are poorly effective. New FA-HNSCC disease models are needed to understand the molecular mechanisms of disease progression, eventually leading to the discovery of drugable targets to delevop new therapies. Here, we describe new cell lines obtained from individual clones of edited CAL27 and CAL33 cells which faithfully phenocopy defective FA pathway, and can be stably grown in standard cultured conditions. We propose these clones as FA-HNSCC disease models that might help to find new therapies.

## Materials and Methods

### Cell culture and genotoxic agents

Tongue SCC cell lines CAL33 and CAL27 were provided by Dr. Silvio Gutkind (UC San Diego, USA). VU1365, a mouth mucosa SCC cell line from a FA patient, was provided by Dr. Josephine Dorsman (Amsterdam UMC, Holland). Clones 34 and 47 from CAL27 (CAL27-c34 and CAL27-c47), clones 5 and 11 from CAL33 (CAL33-c5 and CAL33-c11) were obtained using CRISPR/Cas9 editing (see below). FANCA complemented VU1365-FANCA cells were made from VU1365 upon transduction with a reported retroviral vector carrying the cDNAs for *FANCA* and the neomycin resistance gene (S11FAIN) using two infection cycles of 24 hours (Antonio Casado et al., 2007). Afterwards, transduced cells were selected with G418 at 0.7mg/ml (Calbiochem) for five days and expanded prior to use. All parental and derivative cells were cultured in Dulbecco’s Modified Eagle’s Medium (DMEM) (Gibco, Thermo Fisher) supplemented with 10% fetal bovine serum (FBS) (Gibco, Thermo Fisher) and maintained at 37°C, in an atmosphere of 5% CO2 and 95% humidity. CAL27 and CAL33 parental and cloned cells were authenticated using short tandem repeat (STR) profiling.

### Cell line nucleofection and CRISPR/Cas9 editing within exon 4 of *FANCA* gene

CRISPR/Cas9 genome editing was performed by ribonucleotide-protein (RNP) nucleofection using the 4D-Nucleofector™ System (Lonza), following the electroporation conditions already optimized in the laboratory, which included the use of 15-EW program and the SE Solution (Lonza). Thus, Streptococcus pyogenes Cas9 nuclease (IDT), together with a chemically modified, stable synthetic guide RNA (crRNA) specific for exon 4 of *FANCA* gene linked to tracrRNA (gGM10) (Synthego) were used (Fig S2A). The range of concentrations used was between 0.5-3 μg for Cas9, and 0.7-4 μg for gGM10 guide. Cell viability was analyzed 24 hours later using flow cytometry (BD LSRFortessaTM, BD Biosciences) and DAPI (Roche) labeling, and data analyzed with FlowJo. Exon 4 from *FANCA* gene was subsequently amplified by PCR, using specific primers (Table S2) and the Herculase II fusion DNA polymerase (Agilent). Previous to that, DNA was purified with the DNeasy Blood & Tissue (Qiagen). Editing efficiency was evaluated via nuclease assay using the corresponding Surveyor® Mutation Detection Kit (IDT). DNA fragments resulting from nuclease activity during the Surveyor assay were separated by electrophoresis in 10% TBE-polyacrylamide gels (ThermoFisher). A quantitative measurement of optical density through Image J was performed to quantify efficiency of CRISPR/Cas9 editing. Optical density (OD) intensity was calculated from both edited DNA bands (ed1 and ed2) and non-edited bands (non-ed) applying the following formula: % edition= [(DO_ed1+ DO_ed2 – 2 x DO_background)/ (DO_non_ed+DO_ed1+DO_ed2 – 3 x DO_background)] x 100. Once nucleofection was performed, individual cell clones were seeded separately in 96-well plates using cell sorting (Influx BD™) and amplified with conditioned medium. *FANCA* gene locus was then PCR amplified and sequenced from genomic DNA of selected clones through the Sanger method (StabVida), using the primer P_GM_Forward4 (Table S2). Afterwards, ICE CRISPR Software Analysis Tool (Synthego) was used to quantify and analyze editing results.

### Protein extraction and western blotting

Protein extracts were obtained using a lysis buffer (Hepes 40 mM, Triton-X100 2%, β-glycerophosphate 80 mM, NaCl 200 mM, MgCl_2_ 40 mM, EGTA 20 mM) supplemented with protease and phosphatase inhibitor cocktails (Roche). Proteins were separated in 4-12% Nu-Page Bis-Tris (FANCA analysis) or NuPAGE 3-8% Tris-Acetate (FANCD2) polyacrylamide gels (Invitrogen) and transferred to nitrocellulose membranes (Amersham Biotech) under wet conditions. Membranes were blocked with 5%-non-fat milk in TBS-Tween (Tris-HCl 20 mM, NaCl 137 mM, Tween-20 0.5%) and incubated with the primary antibodies anti-FANCA (1:1000, Abcam), anti-FANCD2 (1:200, Santa Cruz Biotechnology), anti-vinculin (Abcam) and anti-GAPDH (1:5000, Santa Cruz). Peroxidase-coupled, secondary antibodies specific for rabbit IgG (1:5000, Amersham,) and mouse IgG (1:5000, Jackson) were also used. Bands were visualized with the luminescence detection kit Super Signal West Pico Chemiluminscence Substrate (Pierce) according to manufacturer’s instructions, and quantified using the Image Lab 5.2.1 software (BioRad). GAPDH and vinculin proteins were selected as loading controls.

### Mitomycin C (MMC) and cisplatin sensitivity assays

To test the MMC and cisplatin-induced cytotoxicity, clonogenic assays on SCC cells, were performed. Thus, 40.000 cells from CAL27, CAL27-c34, CAL27-c47, VU1365 and VU1365-FANCA, and 100.000 cells from CAL33, CAL33-c5 and CAL33-c11were seeded on 6-well plates and 24 hours later treated with a range of MMC or cisplatin concentrations for 1 hour. Afterwards, cells were washed three times with PBS and cultured in DMEM until control cells reached the maximum confluence (5 days). Finally, cells were stained with crystal violet and eluted with 33% acetic acid. Color intensity was measured using the Genius Pro (Tecan) microplate reader (absorbance at 620 nm). Each experiment was carried out at least three times. The concentrations of MMC and cisplatin corresponding to its 50 inhibitory concentration (IC50) were calculated.

### FANCD2 immunofluorescence

For the FANCD2 immunofluorescence, 40.000 cells from CAL27, CAL27-c34, VU1365 and VU1365-FANCA, or 100.000 cells from CAL33, CAL33-c11 were cultured on 2-well coverslips (Chamber cell culture slides, FALCON) and treated with IC50 concentration values of mitomycin C (MMC) for 1 hour. Subsequently, cells were fixed with 4% formaldehyde for 15 minutes, permeabilized with 0.5%-Triton-X100 for 5 minutes and blocked with 0.1%-NP-40 FBS (Sigma-Aldrich) for 1 hour, to avoid non-specific binding. Thereafter, cells were incubated with the primary antibody anti-FANCD2 (1:200, Abcam) at 4°C for 12 hours. Fluorochrome-complex rabbit IgG specific antibody AlexaFluor 594, (1:1000, Jackson) was used as a secondary antibody. Finally, mounting of slides was carried out using MOWIOL with DAPI (200 μg/ml) for nucleus detection. Visualization was performed using Axioplan 2 imaging (Zeiss) microscope, and images were captured with AxioCam MRm (Zeiss) and visualized through the AxioVision Rel.4.6 (Zeiss) software.

### Chromosome fragility and G2 arrest by the flow cytometry micronucleus (MN) test

Cells were processed by flow cytometry following the procedure previously described by Avlasevich et al (Avlasevich et al., 2006) and reported with some modifications by Hernandez et al (Hernandez et al., 2018). Briefly, 10.000 cells from each cell line were seeded in 96-well plates. The next day cultures were untreated or treated with 0.05 μg/mL of diepoxybutane (DEB) and kept in culture at least one population doubling. Cells were then sequentially stained; first with ethidium monoazyde bromide (EMA) (0.025 mg/mL) and secondly, in lysis solution, with 4’,6-diamidino-2-phenylindole (DAPI) (2 μg/mL). After that, samples were stored at 4°C until being processed in a MACSQuant Analyzer 10 cytometer (Miltenyi). Collected data was analyzed by Flow Jo VX software. The data of micronuclei (MN) and G2 arrest presented in this work represents results from three independent experiments each one in duplicate.

### Cell growth assays

2.000 cells from CAL27, CAL27-c34, CAL27-c47, VU1365 and VU1365-FANCA, and 4.000 cells from CAL33, CAL33-c5 and CAL33-c11were seeded in 96-well plates and cell viability was determined with the colorimetric Cell Proliferation Kit II (XTT) (Roche) 24, 48, and 72 hours later, following the manufacturer’s instructions. Absorbance values were measured using the Genius Pro (Tecan) microplate reader. Each experiment was performed at least three times, with six replicates of each cell line for assay. Background absorbance was subtracted, and data was normalized to the corresponding values of 24 hours measurement. Cells were maintained either in 10% FBS or 0% FBS.

### Wound healing assay for cell migration analysis

Cell migration was analyzed using the *in vitro* wound healing method. Briefly, 400.000 cells from CAL27, CAL27-c34, VU1365 and VU1365-FANCA, or 600.000 cells from CAL33, CAL33-c11 were seeded in 6-well plates and 24 hours later the FBS concentration was reduced to 1% in order to stop cell proliferation. Cells were maintained under these conditions for 72 hours, and after this time, two straight lines were scratched within each plate, creating wounds of approximately 1 mm. Wound closure area was calculated 0, 24, 48 and 72 hours later. Gap distances were quantitatively measured using Image J software. Wound area quantification data are mean ± SEM of three different wounds of three independent experiments.

### Statistical analysis

The half maximal inhibitory concentration (IC50) values are defined as the drug concentrations required for 50% loss of cell viability in a clonogenic assay, using a non-linear regression fit. Data are shown as mean±standar error of the mean (SEM). For cell proliferation and cell migration assays, one-way ANOVA or two-way ANOVA were done respectively. A p-value>0.05(ns), ≤0.05(*), ≤0.01(**), ≤0.001 (***), ≤0.0001 (****) was considered. Graphs and statistics were done using GraphPad Prism 8 software.

## Acknowledgements

We thank Juan Bueren for helpful discussions and critical reading of the manuscript. The authors are also indebted to the patients with FA, their families and clinicians from the Fundación Anemia de Fanconi.

## Author contributions

Conceptualization: R.G.-E., C.S., R.E., J.S., P.R.

Methodology: R.G.-E., C.S., R.E., C.L., M.J.R., F.J.R.-R., J.A.C.

Validation: R.G.-E., C.S., R.E.

Formal analysis: R.G.-E., C.S., R.E., C.L., M.J.R.

Investigation: R.G.-E., C.S., R.E., C.L., P.M., E.S, M.J.R., J.P., J.A.C., E.S.

Resources: H.H., P.R., F.J.R.-R.

Writing – original draft preparation: R.G.-E., C.S., R.E.

Writing – review and editing: R.G.-E., C.S., R.E., J.S., H.H., C.L., M.J.R., P.R.

Visualization: R.G.-E., C.S., R.E.

Supervision: R.G.-E., C.S.

Project administration: R.G.-E., C.S.

Funding acquisition: R.G.-E., C.L., J.S.

## Supplementary Figure Legends

**Figure S1.** Expression values of *FANCA* gene mRNA from The Cancer Cell Line Encyclopedia collection. Cell lines are grouped by tissue location. Expression value positions of CAL27 and CAL33 cells are shown.

**Figure S2. A)** Recognition sequence and cleavage site of guide RNA gGM10 (green) in the exon 4 of the FANCA gene, 3 nucleotides before the PAM-sequence (orange). **B)** Editing analysis of clon 34 from CAL27 (CAL27-c34) after Sanger sequencing. **C)** Editing analysis of clon 47 from CAL27 (CAL27-c47) after Sanger sequencing.

**Figure S3. A)** Editing analysis of clon 5 from CAL33 (CAL33-c5) after Sanger sequencing. **B)** Editing analysis of clon 11 from CAL33 (CAL33-c11) after Sanger sequencing.

